# Histopathology-inferred spatial transcriptomics characterizes the tumor microenvironment in 1,500 head and neck tumors and predicts clinical outcomes

**DOI:** 10.64898/2026.05.16.725687

**Authors:** Sumona Biswas, Sumeet Patiyal, Tien-Hua Chen, Amos Stemmer, Saugato Rahman Dhruba, Sumit Mukherjee, Thomas Cantore, Eldad D. Shulman, Emma Campagnolo, Benjamin H Jenkins, Shyh-Kuan Tai, Pen-Yuan Chu, Ying-Ju Kuo, Yi-Chen Yeh, Chi-Ping Day, Christopher J Hanley, Gareth J Thomas, Muh-Hwa Yang, Danh-Tai Hoang, Eytan Ruppin

## Abstract

Head and neck squamous cell carcinoma (HNSC) is a prevalent malignancy associated with poor prognosis despite recent therapeutic advances. We hypothesized that a comprehensive understanding of the spatial heterogeneity and organization of the tumor microenvironment (TME) can substantially improve risk stratification and prediction of treatment response in HNSC. As spatial transcriptomics (ST) remains labor-intensive and costly, we developed *HEiST* (H&E-Inferred Spatial Transcriptomics), a deep learning framework that predicts spatially resolved gene expression profiles directly from routine hematoxylin and eosin (H&E)-stained histology slides. After rigorous validation across two independent external ST cohorts, we applied *HEiST* to infer spatial transcriptomes across 1,500 HNSC patient tumors spanning two publicly available datasets and two newly generated cohorts, one treated with concurrent chemoradiotherapy (CCRT) and one with immunotherapy. This large-scale analysis uncovered reproducible spatial clusters characterizing the HNSC TME, defining two distinct prognostic *Spatiotypes*, *Immune*-*Exhausted* and *Immune*-*Activated*, with significantly distinct survival outcomes. Critically, spatial cluster composition accurately predicts HPV status and yields treatment response predictors for both CCRT/radiotherapy and immunotherapy that outperform costly gene-expression and direct image-based approaches. Notably, the ST cluster–based predictor of immunotherapy response markedly surpasses the performance of commonly used FDA-approved biomarkers, including CPS, TPS, and their combination. To the best of our knowledge, this represents the first virtual spatial profiling effort and the most comprehensive large-scale spatial TME analysis in HNSC to date. *HEiST* thus introduces a scalable, low-cost, and spatially grounded biomarker discovery for precision oncology in HNSC.

## Introduction

Head and neck squamous cell carcinoma (HNSC) comprises a heterogeneous group of malignancies originating from the epithelial lining of anatomically and functionally intricate sites, including oral cavity, pharynx, and larynx. HNSC accounts for ∼4–5% of the global cancer burden, with over 900,000 new cases diagnosed annually [1, 2]. Biologically, HNSC is broadly divided into HPV-positive tumors driven by high-risk HPV infection and HPV-negative tumors, typically associated with tobacco and alcohol exposure [3]. These subgroups differ fundamentally in their mutational landscape, immune microenvironment, therapeutic responsiveness, and clinical outcome. Integrative genomic and transcriptomic analyses have further identified reproducible molecular subtypes in HNSC (basal, classical, mesenchymal, and HPV-driven), each characterized by distinct gene expression profiles, genomic alterations, and variable stromal and immune enrichment [4]. Despite these advances, clinical risk stratification and treatment selection in clinical practice remain largely anchored in conventional H&E-stained histopathology sections, including assessment of tumor grade, invasive growth patterns, tumor budding and stromal architecture [5, 6, 7, 8]. This framework is supplemented by a limited repertoire of immunohistochemical biomarkers, principally p16 for HPV status [9] and PD-L1 expression for anti-PD1 checkpoint immunotherapy eligibility [10]. Notably, established molecular classifications are derived from bulk profiling, which averages signals across heterogeneous tumor, stromal, and immune compartments and is devoid of spatial relationships within the tumor microenvironment (TME). In contrast, H&E sections capture rich morphological information that reflects underlying biological states and carries independent prognostic values. However, the extent to which H&E images encode gene expression programs correlated with clinically actionable risk remains largely unexplored. Here, we address this challenge by studying whether and to what extent routinely available H&E slides can be utilized to comprehensively characterize the HNSC TME and identify accessible, scalable biomarkers for clinical outcomes stratification.

Spatial transcriptomics (ST) technology has emerged as a powerful tool that enables unprecedented insights into tissue organization, cellular characteristics, functions, and interactions within the TME while preserving the spatial context of cells. Recent ST studies in HNSC have already yielded mechanistically and clinically significant discoveries. For instance, the spatial co-localization of fibroblastic reticular cell (FRC)-like with CD4⁺ T cells and B cells has been shown superior outcomes prediction in patients receiving immune checkpoint inhibitors [11], while epithelial–mesenchymal transition (EMT)-driven tumor evolution and resistance to anti-HER therapy have been linked to MET/PDGF signaling [12]. In another study in oral squamous cell carcinoma (OSCC), cancer-associated fibroblast (CAF)-secreted TGF-β1 has been demonstrated to promote metastasis via EMT [13]. Furthermore, ST investigations have characterized distinct transcriptional programs between the tumor core and invasive leading edge [14], and identified ferroptosis–immune colocalization at the invasive front as a determinant of immune checkpoint blockade sensitivity [15]. Subsequently, additional studies have nominated PKM2 as a stemness regulator and EPHA2 as an angiogenesis driver in HPV-negative disease [16], and have revealed metabolic heterogeneity as a key architect of immunosuppression within the TME [17]. Collectively, these findings highlight the translational potential of ST technology in unraveling the complex spatial architecture of the TME, identifying therapeutic targets and uncovering predictive biomarkers to guide treatment in HNSC. Despite these advances, the widespread clinical adoption of ST remains limited by high cost, technical complexity, and tissue requirements that preclude its routine deployment in large-scale clinical cohorts. As a result, spatially resolved transcriptomic data are unavailable for most HNSC patients, thereby limiting novel population-level biomarker discovery, predictive modeling of therapeutic response, and the broader applicability of precision oncology to real-world clinical practice.

To overcome these limitations, several deep learning methods have been developed to infer spatial gene expressions directly from routinely available histopathology images. These methods utilize a wide range of deep learning architectures including convolutional neural networks (STNet [18], BrST-Net [19], DeepSpaCE [20]), attention models (HisToGene [21], CarHE[22]), capsule networks (THItoGene [23]), vision transformer-GNN hybrid model (FmH2ST [24], Hist2ST [25]), as well as contrastive learning approaches (BLEEP [26], Reg2ST [27], mclSTExp [28]), and interpretable frameworks (Stimage [29]). Indeed, such progress demonstrates the feasibility of learning complex relationships between tissue morphology and ST profiles. However, existing approaches have largely been developed and evaluated on limited datasets and selected cancer types, most notably breast [31, 32] and colorectal cancers [32], leaving the spatial complexity and clinical heterogeneity of HNSC largely unaddressed.

Given this gap, we investigated whether deep learning–based spatial inference from routine histology can robustly capture biologically and clinically relevant spatial programs in HNSC across large and diverse patient cohorts— an area that remains largely unexplored. Building on our recently developed *Path2Space* model [30], we tailored an HNSC-specific ST inference model termed *HEiST* (H&E-inferred Spatial Transcriptomics) to predict spatial gene expression from H&E images. *HEiST* was trained and rigorously cross-validated on an HNSC ST patient cohort, then validated across two independent external HNSC ST cohorts, establishing a set of approximately 2,800 genes whose spatial expression are reliably inferred from routine H&E slides. We pursued two main translational objectives: (i) characterizing the spatial organization of the HNSC TME at scale, and (ii) identifying spatially resolved predictors of patient survival and treatment response.

Applying *HEiST* to predict the spatial gene expression across a large cohort from The Cancer Genome Atlas (TCGA) [33], we identified 11 shared spatial TME clusters, whose composition across tumors revealed two previously unrecognized HNSC prognostic patient subgroups, termed *Spatiotypes*. We independently validated the prognostic relevance of *SpatioTypes* on two large external HNSC cohorts, HANCOCK [34] and a newly generated Taiwan CCRT. Beyond prognosis, the spatial clusters demonstrated strong discriminative power for HPV status stratification, outperforming direct H&E image-based approaches. Applying *HEiST* to two large treatment cohorts (one cohort treated with concurrent chemoradiotherapy (CCRT)/radiotherapy [33], and another with immune checkpoint blockade (ICB) therapy [prospectively collected and unpublished]), we discovered spatially defined biomarkers associated with therapeutic response. These spatial features significantly outperform measured transcriptomics, slide-inferred transcriptomics, and direct H&E image-based models in predicting patients’ treatment response. Furthermore, these spatial biomarkers surpassed FDA-approved clinical biomarkers (CPS, TPS, and their combination) in stratifying ICB-treated patients. To the best of our knowledge, this study presents the first scalable HNSC-specific framework for virtual spatial profiling derived directly from routine histology, and the most comprehensive spatial characterization of the HNSC TME to date. Collectively, *HEiST* paves the way for spatially informed precision oncology in HNSC, enabling spatially grounded biomarker discovery and predictive modeling of treatment response at scale.

## Results

### Study overview

Our study was performed in two major steps: (1) We first developed a deep learning framework, *HEiST*, to predict spatial gene expression directly from routine H&E-stained tissue sections in HNSC, utilizing ST datasets from HNSC patient cohorts. (2) We then applied the trained *HEiST* predictor to large patient cohorts where only H&E images were available (i.e., without ST measurements), enabling large-scale deep learning-based spatial inference of TME organization. These large-scale spatial predictions enable the identification and validation of spatial biomarkers associated with patient outcomes (**Fig. 1**).

**Fig. 1.**
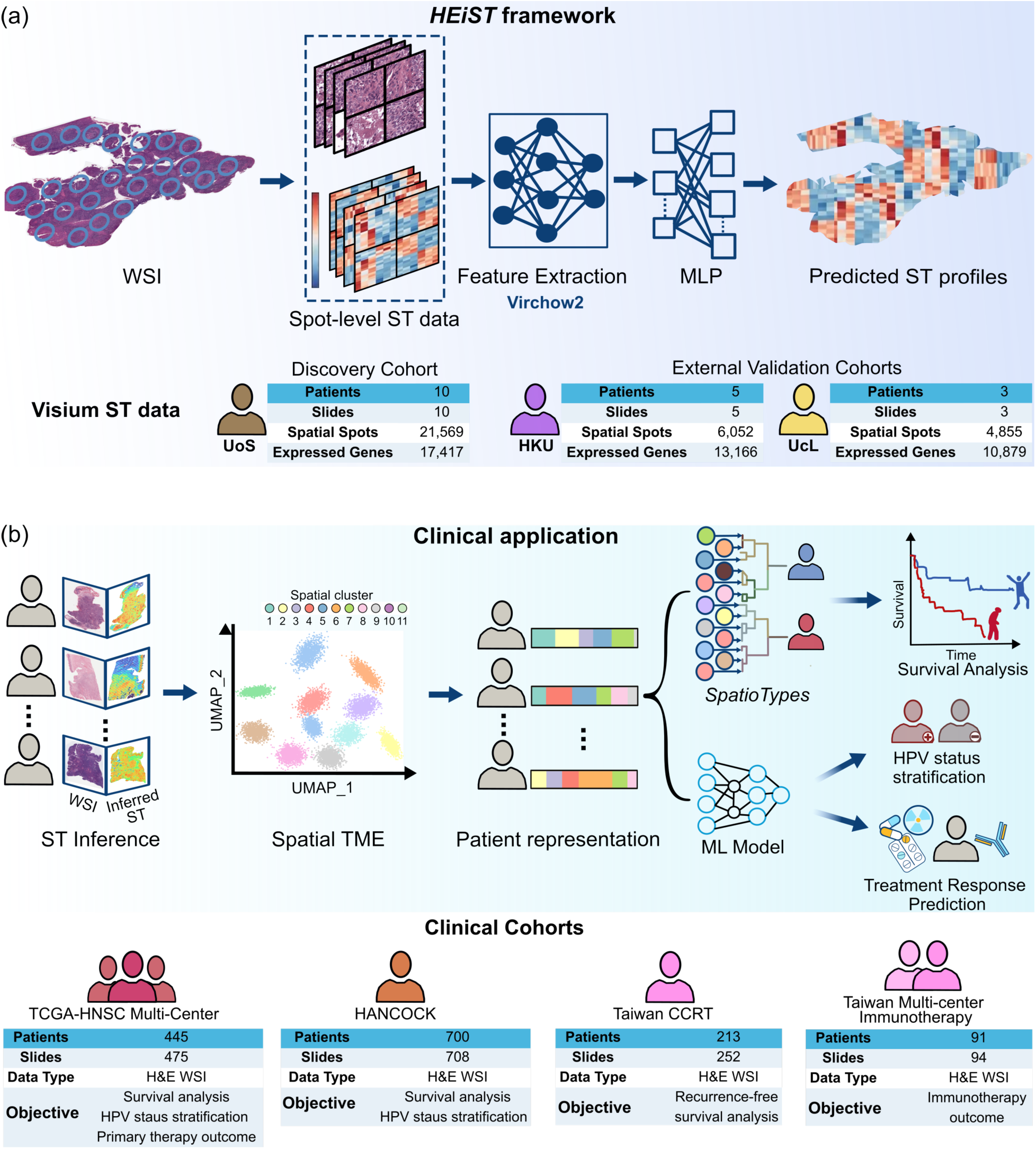
Overview of *HEiST* framework and clinical application. **(a)** H&E inferred Spatial Transcriptomic (*HEiST*) framework. *HEiST* was trained on paired spot-level image tiles and spatially resolved gene expression. The discovery cohort comprises samples from the newly generated University of Southampton (UoS; 21,569 spatial spots, 17,417 expressed genes; 10 patients, 10 whole slide image (WSI) slides) cohort. The external validation cohorts include the University of Hong Kong (HKU; 6,052 spatial spots, 13,166 expressed genes; 5 patients, 5 slides) and UCLouvain (UcL; 4,855 spatial spots, 10,879 expressed genes; 3 patients, 3 slides). After quality control and color normalization, image tiles were processed through the pathology foundation model Virchow2 [35] for feature extraction and then fed into multilayer perceptron (MLP) layers to predict spot-level gene expression for 17,417 genes. **(b)** Spatial analyses across clinical cohorts. During inference, H&E tiles undergo the same preprocessing steps and were used as input to infer spatial gene expression. The inferred expression data was utilized to identify 11 spatial tumor microenvironment (TME) profiles. These profiles were leveraged in two complementary ways. First, an unsupervised clustering algorithm was applied to stratify tumors into distinct, prognostic, spatially grounded subgroups, termed *SpatioTypes*, within the TCGA-HNSC cohort. The prognostic value of these *SpatioTypes* was externally validated in two independent clinical cohorts, HANCOCK, and freshly collected from the Taiwan CCRT cohort. Second, three supervised machine learning models, each independently trained on the 11 spatial TME clusters to predict (i) HPV status assessed in the TCGA-HNSC and HANCOCK cohort, (ii) response to concurrent chemoradiotherapy/radiotherapy examined in the TCGA-HNSC cohort, and (iii) response to immunotherapy evaluated in the prospectively collected Taiwan Multicenter Immunotherapy cohort.

In the first step, following the approach we developed and established in *Path2Space* [30], our histology image-based ST prediction framework, *HEiST* leverages a recent pathology foundation model, Virchow2 [35] as an image feature extraction module, followed by a multilayer perceptron (MLP) regressor to predict gene expression profiles (**Fig. 1a**). *HEiST* was trained using 10x Genomics Visium ST data, which quantifies gene expression at spots with a 55-µm diameter (∼100-µm center-to-center spacing) spatially registered to the corresponding whole slide image (WSI). For each Visium spot, image tiles were extracted from the spatially corresponding H&E region and subject to preprocessing, including quality check and color normalization, prior to the feature extraction (see **Methods** for more details). The Virchow2-based feature extraction module generates high-dimensional image representations from the preprocessed H&E-stained tiles, which were then used as input to the MLP regressor to predict a spot-level gene expression vector of 17,417 genes (genes detected in ≥5% of spots on each slide).

*HEiST* was trained on the newly generated University of Southampton (UoS) HNSC ST discovery cohort [11] comprising 21,466 Visium spots from 10 patients (10 WSI slides), which enables the model to learn predictive image–expression relationships under spot-level supervision. Subsequently, we evaluated the robustness and generalizability of the model on two independent HNSC ST validation cohorts, comprising a total of 11,485 additional Visium spots from 8 unseen slides. These included 6,097 spots from 5 patients from the University of HongKong (HKU) cohort [13] and 5,388 spots from 3 patients from the UCLouvain (UcL) cohort [12].

In the second step, we applied the trained *HEiST* model to four large independent HNSC patient cohorts, including TCGA-HNSC, HANCOCK, and two prospectively collected Taiwan cohorts comprising approximately 1,500 patients to identify and characterize spatial TME architecture associated with survival. Finally, we utilized these spatial TME profiles to discover robust, clinically relevant biomarkers predictive of HPV status, validated across two independent patient cohorts, and treatment response, confirmed across two cohorts receiving distinct therapeutic regimens (**Fig. 1b**). Overall, the spatial biomarker–based model demonstrated superior predictive performance compared to the established biomarker-based models.

### *HEiST* robustly predicts the expression of nearly 3,000 genes at spatial resolution

The performance of the *HEiST* was first assessed using leave-one-patient-out cross-validation scheme on the UoS discovery cohort, in which the model was trained on all patients except one and evaluated predictions on the held-out patient’s slides (See **Methods** for details). Subsequently, gene-wise prediction accuracy was quantified using the Pearson correlation coefficient (PCC) between ground-truth and predicted expression values across all spots in each slide. The model achieved an overall median PCC of 0.48, with 10,162 robustly predicted genes (defined by PCC>0.4, out of 17,417 genes, **Fig. 2a**). We then evaluated the model generalizability on H&E whole-slide images (WSIs) from two independent external ST cohorts, HKU (5 patients, 6,052 spots) and UcL (3 patients, 4,855 spots). *HEiST* attained a median PCC of 0.24, with 2,831 robustly predicted genes (out of 13,751 genes, **Fig. 2b**). Notably, 2,772 genes showed consistent, robust prediction (PCC>0.4) across all three (cross-validation and both external validation) cohorts (**Fig. 2c**).

**Fig. 2.**
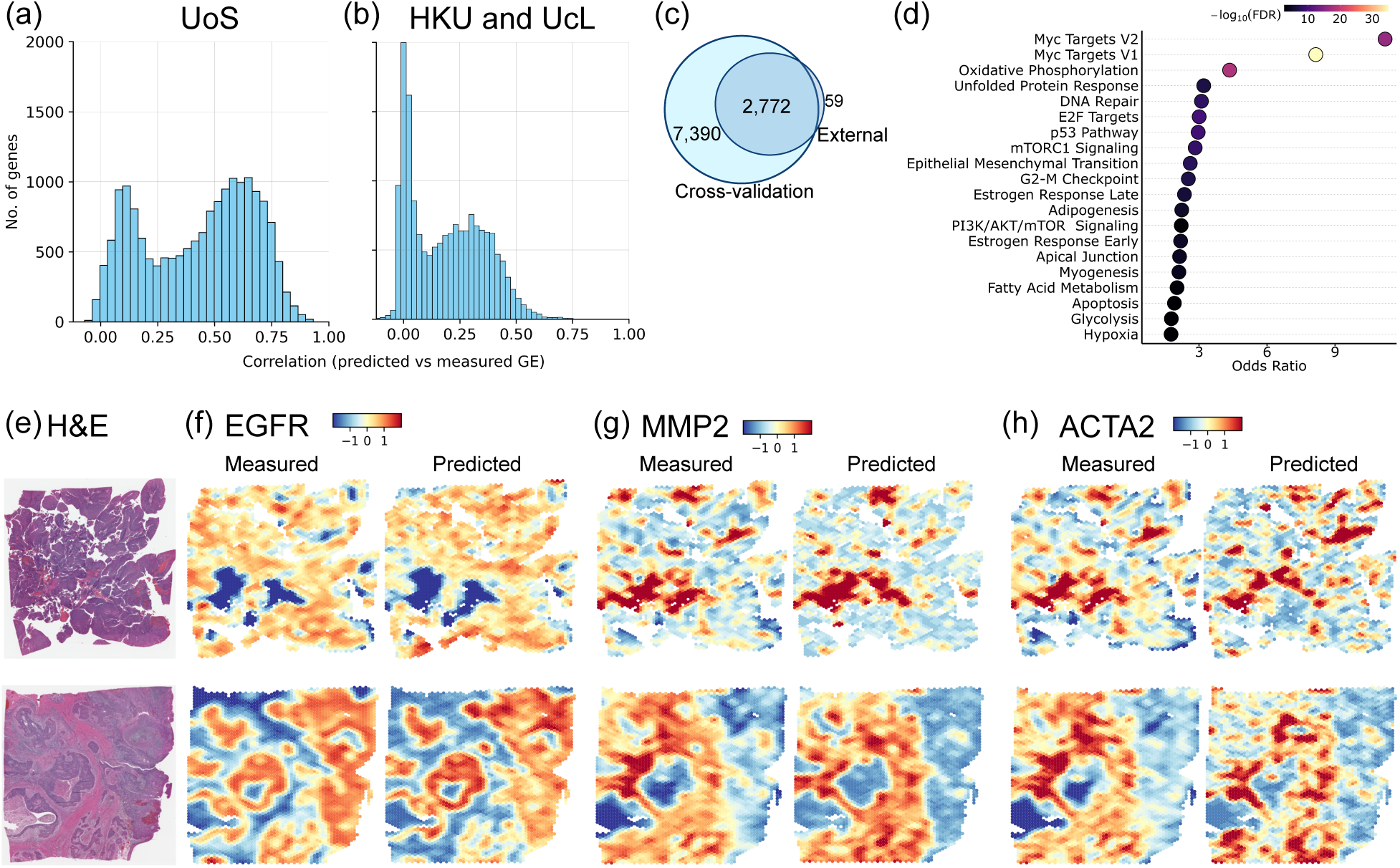
*HEiST* performance in predicting spatial gene expression. (**a, b**) Performance of spatial gene expression prediction is shown as distributions of Pearson correlation coefficients (PCCs). Results are shown for discovery (in cross-validation) on the UoS cohort (panel **a**) and independent validation on HKU and UcL cohorts (panel **b**). (**c**) Venn diagram of robustly predicted genes (PCC> 0.4) between discovery and combined validation cohorts. (**d**) MSigDB Cancer Hallmark pathway enrichment for 2,772 robustly predicted genes across three cohorts, with enrichment strength shown as odds ratios and significance indicated by −log10 adjusted *P* values (adjusted *P*≤0.05). (**e–h**) Representative examples from the discovery cohort, including H&E images (**e**) and spatial expression maps of three robustly predicted genes (relevant to HNSC biology and prognosis), EGFR (f), MMP2 (**g**), and ACTA2 (**h**), showing measured (ground-truth) and *HEiST-*predicted expression across tissue sections. Panels (**f–h**) show scaled spatial expression for two representative cross-validation slides per gene, with measured expression displayed on the left and *HEiST*-predicted expression on the right.

To learn more about the biological relevance of these 2,772 robustly predicted genes, we performed gene set enrichment analysis (GSEA) using 50 cancer hallmarks [36]. This analysis revealed significant enrichment across multiple hallmark cancer pathways (**Fig. 2d**), most prominently MYC targets (V1/V2), oxidative phosphorylation, unfolded protein response, DNA repair, E2F, and p53 signaling, along with additional oncogenic and TME–associated programs such as epithelial–mesenchymal transition, mTORC1, adipogenesis, and PI3K/AKT/mTOR signaling. Notably, the robustly predicted gene set includes EGFR, MMP2, and ACTA2, all of which have established relevance to HNSC biology and prognosis, as shown in **Fig. 2f–h**. EGFR drives proliferation and therapeutic resistance [37], MMP2 promotes tumor invasion through extracellular matrix degradation [38], and ACTA2 marks cancer-associated fibroblasts linked to stromal remodeling and poor outcomes [39]. Together, these findings underscore that the *HEiST* not only generalizes well across independent validation cohorts generated under different experimental settings but also preferentially captures gene expression patterns from H&E morphology linked to core cancer hallmarks and key regulatory pathways in HNSC.

### The TCGA-HNSC cohort spans two biologically distinct tumor *SpatioTypes*

Having established the predictive power of *HEiST*, we next applied it to the TCGA-HNSC cohort (475 H&E-stained slides, 445 patients) to infer spatial gene expression across 2,772 robustly predicted genes and systematically characterize TME spatial organization at the population scale.

To this end, we divided each TCGA-HNSC slide into a grid of approximately 10,000 “pseudo-spots” per slide, matching the spot dimensions of Visium training dataset, but forming a contiguous manifold without any inter-spot gaps. Subsequently, to identify transcriptionally and spatially consistent tissue domains across all samples in the TCGA-HNSC cohort, we applied a well-established graph convolutional framework SpaGCN [40] which integrates the predicted ST profiles with pseudo spatial coordinates and spatial adjacency to detect spatial domains within each sample (**Fig. 3a**). To ensure consistent clustering annotation across all slides, we performed joint clustering based on spatial domain-averaged expression and identified 11 reproducible spatial clusters that encompass the detected spatial domains, as visualized by UMAP displayed in **Fig. 3b**. Each SpaGCN-identified domain was then assigned to one of these 11 reference clusters (**Fig. 3c**) (See **Methods** for details). The GSEA of these 11 spatial clusters captures shared HNSC microenvironmental structures (**Fig. 3d, e,** and **Supplementary Fig. 2a**) whose relative abundances vary among the individual samples in the cohort. Indeed, some tumors are dominated by immune-active niches while others are enriched with immune-suppressed spatial clusters with high proliferation, suggesting that their distinct spatial organization may underlie variability in clinical outcomes across HNSC patients. We thus hypothesized that a tumor’s spatial cluster abundance could serve as a promising spatially grounded biomarker in HNSC.

**Fig. 3.**
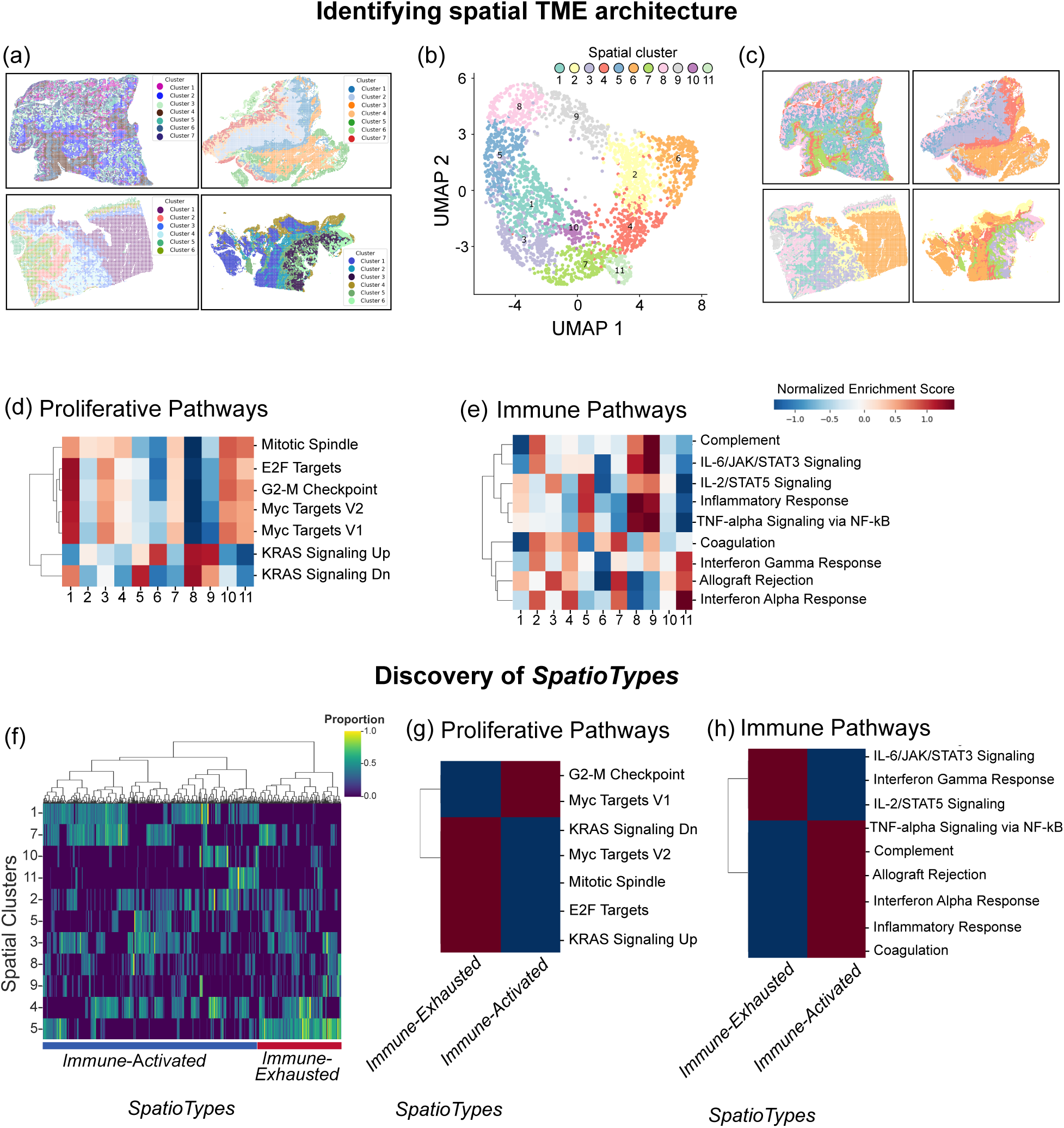
Discovery of two distinct TME *SpatioTypes*. (**a**) Example slides displaying SpaGCN-identified spatial domains in individual TCGA-HNSC samples. (**b**) Uniform Manifold Approximation and Projection (UMAP) visualization of 11 spatial clusters identified in the TCGA-HNSC cohort. (**c**) Representative TCGA-HNSC slides showing spatial cluster assignments across the 11 shared clusters. (**d, e**) Pathway-level activity across spatial clusters is shown as heatmaps of normalized enrichment scores for proliferative (panel **d**) and immune-related (panel **e**) pathways. Hallmark gene sets are from MSigDB (metabolic and stress, see **Supplementary** Fig. 2a). (**f**) Unsupervised hierarchical clustering of TCGA-HNSC samples (n=445) based on the proportions of the 11 spatial clusters stratifies patients into two *SpatioTypes*. Heatmap rows correspond to spatial clusters and columns to patients, with values indicating cluster proportions. The *Immune-Exhausted* and *Immune-Activated SpatioTypes* are highlighted by the color bar at the bottom. (**g, h**) Aggregated MSigDB Hallmark pathway activity across tumor *SpatioTypes*. Proliferative (panel **g**) and immune (panel **h**) pathways display distinct enrichment patterns between *Immune-Exhausted* and *Immune*-*Activated SpatioTypes*. See **Supplementary** Fig. 2c for metabolic and stress adaptive pathway enrichment analysis.

We next characterized each sample by its spatial cluster abundance and applied hierarchical clustering across the entire TCGA-HNSC cohort to understand about their possible prognostic relevance (**Fig. 3f**). Interestingly, this analysis consistently demonstrated two optimal patient-level subgroups, termed *SpatioTypes,* across multiple metrics, including silhouette score, Davies-Bouldin index, and Calinski-Harabasz index (**Supplementary Fig. 2b**). These subgroups display distinct hallmark pathway activities (**Fig. 3g, h**) and TME architectures. The *Immune-Exhausted SpatioType* exhibits dysfunctional immune responses, proliferation, and stress-adaptive metabolic reprogramming, whereas the *Immune-Activated SpatioType* demonstrates coordinated immune engagement with suppressed proliferation and metabolic differentiation ( **Supplementary Fig. 2c**).

### The two major HNSC *SpatioTypes* are associated with distinct patterns of patient survival

To quantitatively evaluate the predictive power of inferred TME compositions in stratifying patient survival, we performed survival analysis using the two *SpatioTypes*. In the TCGA-HNSC cohort, we found that *Immune-Exhausted* patients exhibited markedly worse survival across all endpoints compared to *Immune*-*Activated* patients, after adjusting for age and tumor stage (OS: HR=1.35, p=0.049; PFS: HR=1.37, p=0.052; DSS: HR=1.43, p=0.059) (**Fig. 4a–c**). This finding is biologically consistent, as the *Immune-Exhausted SpatioType* represents a chronically inflamed yet dysfunctional TME, where immune exhaustion and cellular stress create a treatment-resistant, pro-tumorigenic microenvironment despite active immune signaling. In contrast, the *Immune*-*Activated SpatioType* reflects an immune-permissive “hot” microenvironment that supports effective anti-tumor immunity. Interestingly, this pattern emerged within the HPV-negative patient subset, where the *Immune-Exhausted SpatioTypes* consistently conferred increased mortality risk (OS: HR=1.27; PFS: HR=1.32; DSS: HR=1.37) though with modest statistical significance (**Fig. 4d–f**). Notably, this analysis was not performed in HPV-positive patients due to severe class imbalance (1 *Immune*-*Exhausted* vs. 41 *Immune*-*Activated*). Reassuringly, most HPV-positive patients belonging to the *Immune*-*Activated Spatiotype* (**Supplementary Fig. 3d**) are biologically consistent with the well-established immune-permissive nature of HPV-positive HNSC.

**Fig. 4.**
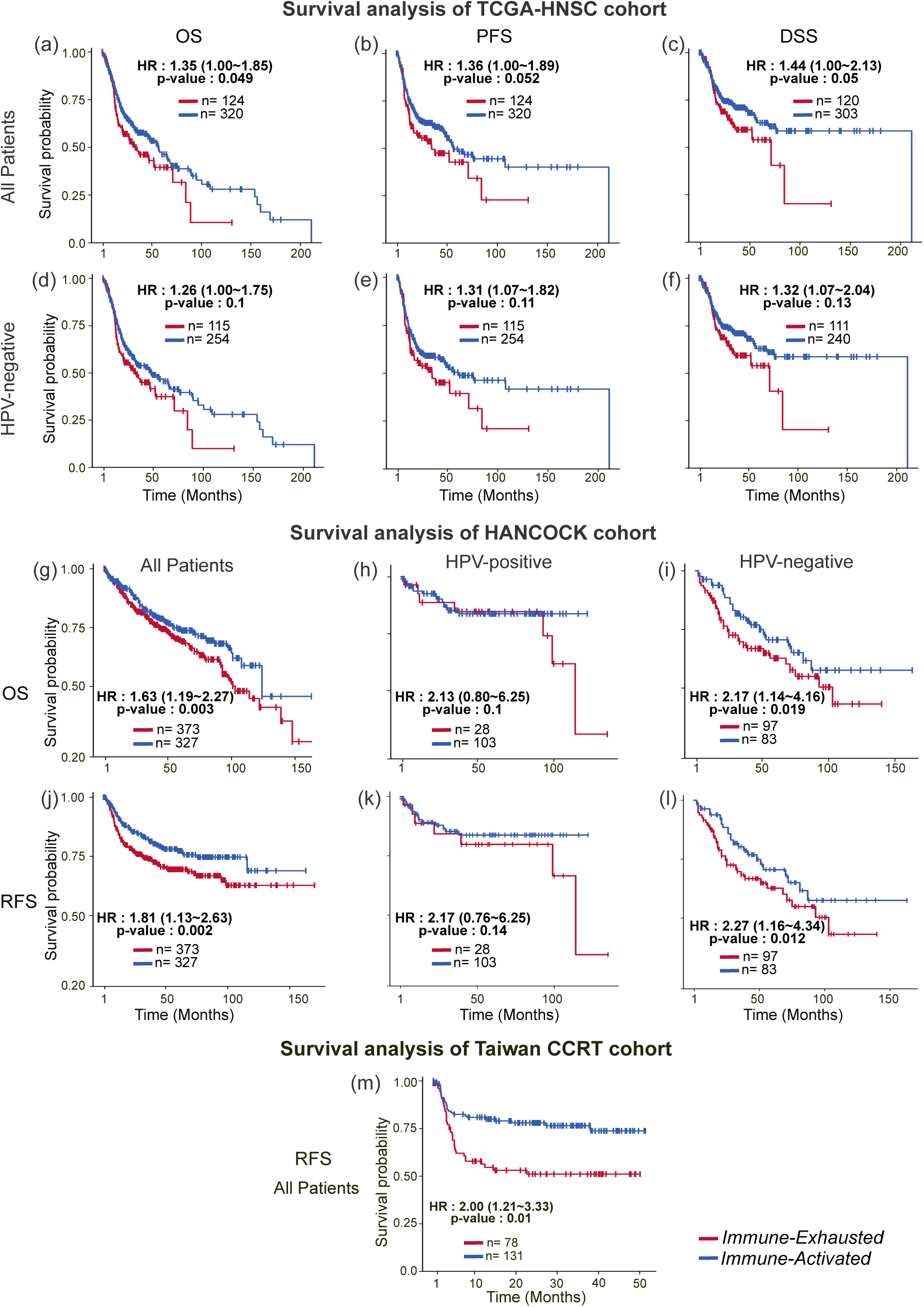
Prognostic value of *SpatioTypes* across three large patient cohorts. (a–c) Kaplan–Meier survival analyses of the TCGA-HNSC cohort comparing *SpatioTypes* across overall survival (OS; panel **a**), progression-free survival (PFS; panel **b)**, and disease-specific survival (DSS; panel **c**). *Immune-Exhausted* patients exhibit significantly worse survival across all endpoints compared to *Immune-Activated* patients (log-rank test, p<0.05 for all endpoints). **(d–f)** Similar to (**a**–**c**) but for the subset of HPV-negative patients. Survival analyses show a consistent trend toward worse OS, PFS, and DFS in the *Immune-Exhausted SpatioType* with modest statistical significance. **(g–l)** Prognostic value of *SpatioTypes* externally validated in the HANCOCK cohort recapitulates the same survival trends observed in TCGA-HNSC across all patients (**g**), HPV-positive (**h**), and HPV-negative (**i**) subgroups for OS and Recurrence-free survival (RFS; **j, k, l**), respectively. (**m**) RFS analysis show consistent *SpatioType*-associated trends in newly collected Taiwan CCRT cohorts.

The prognostic value of the *SpatioTypes* was further examined in a large, publicly available independent validation cohort, HANCOCK (708 slides from 700 patients treated with systemic therapy, radiotherapy, or chemoradiotherapy). Following the same spatial inference pipeline used for the TCGA-HNSC cohort, we inferred ST profiles from the HANCOCK slides and transferred the 11 TCGA-HNSC-derived spatial cluster labels to this dataset (see **Methods** for details). The survival analysis further validated the *SpatioType* stratification. Showing consistent stratification for both OS and RFS after adjusting for age and tumor stage (OS: HR=1.63, p=0.003; RFS: HR=1.81, p=0.002; **Fig. 4g, j**). The hazard ratio remained consistently higher in *Immune*-*Exhausted* than *Immune*-*Activated SpatioType* across HPV status, with modest effect in HPV-positive patients (OS: HR=2.13, p=0.1; RFS: HR=2.17, p=0.14; **Fig. 4h, k**), likely reflecting the predominantly *Immune*-*Activated* biology of HPV-positive tumors (102 *Immune*-*Activated* vs. 28 *Immune*-*Exhausted*) consistent with observations in TCGA-HNSC. In contrast, statistically significant associations were observed in HPV-negative patients (OS: HR=2.17, p=0.019; RFS: HR=2.2, p=0.012; **Fig. 4i, l**).

Next, we tested the robustness of *SpatioTypes*-based stratification in a new, prospectively collected cohort of HNSC patients treated with concurrent chemoradiotherapy (CCRT), Taiwan CCRT cohort (252 H&E-stained slides, 213 patients). Applying our spatial inference pipeline and TCGA-HNSC cluster transfer, patient-level clustering successfully reproduced the prognostic stratification observed in TCGA-HNSC and HANCOCK cohorts. Specifically, *SpatioTypes* significantly predicted recurrence-free survival (RFS) after adjusting for age and stage (RFS: HR=2.08, p=0.01; **Fig. 4m**). In summary, this cross-cohort validation across ∼1,400 patients demonstrates that *SpatioTypes* derived from inferred spatial expression capture clinically relevant tumor biology driving disease progression.

### The Spatial TME architecture is conserved across TCGA-HNSC, HANCOCK, and Taiwan CCRT

We next sought to determine whether the biological TME programs defining each *SpatioType* were preserved across TCGA-HNSC, HANCOCK, and Taiwan CCRT cohorts. To this end, we compared the MSigDB Hallmark pathway enrichment patterns across the cohorts. Remarkably, the normalized enrichment scores across four functional categories— proliferation, immune, metabolism, and stress-adaptive revealed highly consistent patterns across all 11 spatial clusters in the three cohorts, with Spearman correlations ranging from 0.96 to 0.98 (**Figs. 5a–e**). Furthermore, principal component analysis (PCA) demonstrated a strong concordance in the overall pathway landscape across the cohorts (**Fig. 5f** for TCGA-HNSC and HANCOCK; **Fig. 5g** for TCGA-HNSC and Taiwan CCRT). In addition, cross-cohort single-sample GSEA confirmed that all pathway activity scores were perfectly correlated across the cohorts, with a Spearman correlation of 0.97 between TCGA-HNSC and HANCOK (**Fig. 5h**), and 0.98 between TCGA-HNSC and Taiwan CCRT (**Fig. 5i**). Collectively, these findings highlight that *HEiST* robustly captures conserved spatial transcriptional programs across geographically and ethnically distinct HNSC populations.

**Fig. 5.**
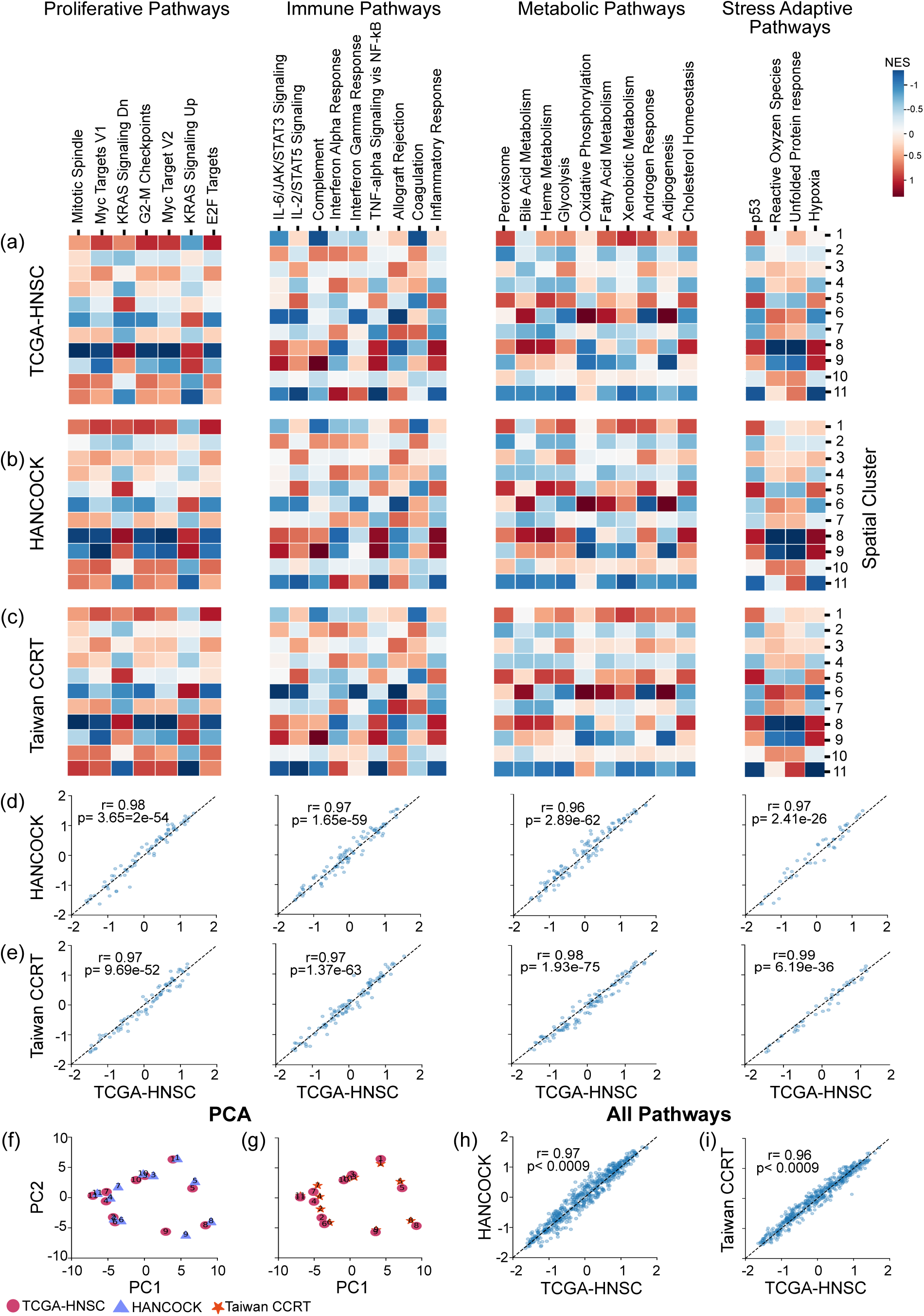
Spatial cluster characteristics are conserved across TCGA-HNSC, HANCOCK, and Taiwan CCRT cohorts. (**a–c**) Heatmaps showing normalized pathway enrichment scores for MSigDB Hallmark pathways across 11 spatial clusters in the TCGA-HNSC (**a**), HANCOCK (**b**), and Taiwan CCRT (**c**) cohorts. Pathways are organized into four functional categories: proliferative (first), immune (second), metabolic (third), and stress-adaptive (fourth). Columns represent Hallmark pathways, and rows represent spatial clusters (1–11). Red and blue indicate positive and negative pathway enrichment scores, respectively (color scale: −1.0 to 1.0). (**d, e**) Spearman rank correlations of normalized pathway enrichment scores between TCGA-HNSC and HANCOCK (**d**), and between TCGA-HNSC and Taiwan CCRT (**e**), demonstrating strong concordance across all four functional pathway categories (Spearman correlation r=0.96–0.98). (**f–i**) Principal component analysis (PCA) of normalized enrichment scores (NES) per spatial cluster in the TCGA-HNSC (circles) and HANCOCK (triangles) cohorts (**f**), and in the TCGA-HNSC (circles) and Taiwan CCRT (stars) cohorts (**g**). Numbers denote spatial clusters 1–11. Overall correlation of normalized enrichment scores between TCGA-HNSC and HANCOCK (**h**), and between TCGA-HNSC and Taiwan CCRT (**i**), across all pathways and clusters (Spearman r=0.96-0.97, p<0.0009).

### The spatial TME composition robustly stratifies HPV status in HNSC

To evaluate the diagnostic utility of the spatial TME for predicting HPV status, we represented each slide as an 11-dimensional vector capturing the fractional abundance of each spatial cluster across the tissue. These features were then used as input to a Logistic Regression model to predict HPV status. Using a five-fold cross-validation within the TCGA-HNSC cohort, the spatial cluster–based model achieved an AUC of 0.88, comparable to two alternative approaches: a model based on measured gene-expression (AUC=0.92) and a model that predicts HPV status from H&E slides without intermediate gene-expression inference (AUC=0.85) (**Fig 6a**). However, when evaluating in external validation, HANCOCK cohort, the inferred spatial cluster–based model achieved an AUC of 0.80, markedly outperforming the direct H&E-based model, which yielded an almost random AUC of 0.52 (**Fig. 6b**). Notably, measured bulk transcriptomics data were not available for the HANCOCK cohort. These findings demonstrate that spatially resolved TME composition captures transcriptional signals predictive of HPV status and provides improved generalizability beyond conventional histological approaches.

**Fig. 6.**
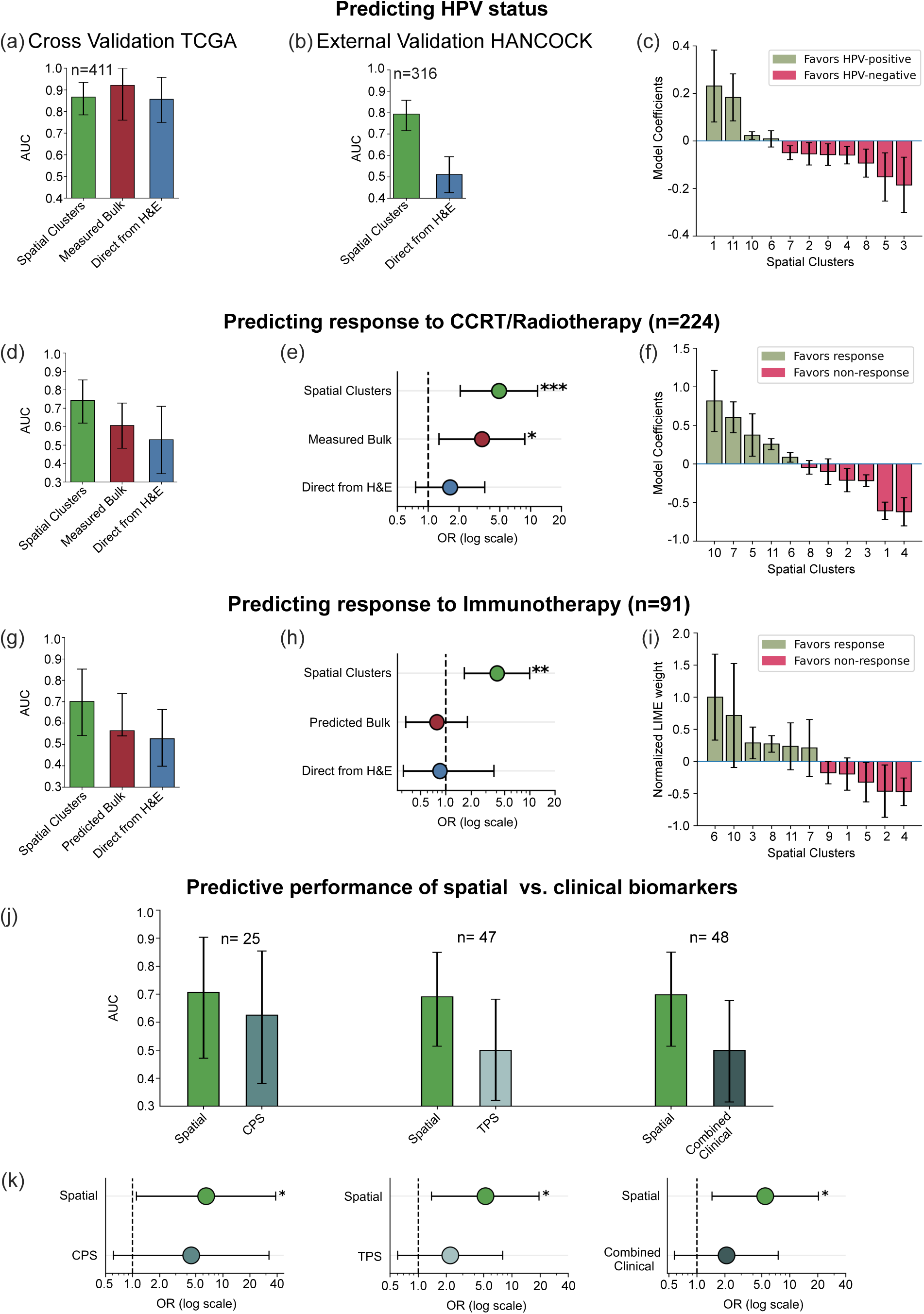
Spatial TME composition from inferred spatial transcriptomics predicts HPV status, and patient response to concurrent chemoradiotherapy/ radiotherapy and immunotherapy. (**a–c**) Prediction of HPV status. The AUC values from cross-validation in the TCGA-HNSC cohort (n=411) (**a**) and external validation in the HANCOCK cohort (n=316) (**b**) achieved by our 11-spatial-cluster-based model (green) are shown in comparison with two traditional approaches: a measured bulk gene expression–based model (red) and a direct pathology image–based model without an intermediate spatial gene expression inference step (blue). Notably, measured bulk expression data were not available for the HANCOCK dataset. (**c**) Contributions of spatial clusters to the prediction of HPV status are shown as logistic regression coefficients. Values are averaged over five-fold cross-validation with standard errors. Positive (mint green) and negative (pink) values indicate associations with HPV-positive and HPV-negative status, respectively. **(d–f)** Prediction of patient response to concurrent chemoradiotherapy/ radiotherapy in TCGA-HNSC cohort (n=224). The AUC value (**d**) and odds ratio (**e**) achieved by our 11-spatial-cluster-based model (green) are shown, in comparison with two traditional approaches: bulk gene expression–based model (red), and a direct pathology image–based model without an intermediate spatial gene expression inference step (blue). (**f**) Contribution of spatial clusters to the prediction of concurrent chemoradiotherapy/radiotherapy is shown as logistic regression coefficients. Values are averaged over five-fold cross-validation with standard errors. Positive (mint green) and negative (pink) values indicate associations with response and non-response, respectively. **(g–i)** Similar to (**d–f**) but for ICB in Taiwan multicenter immunotherapy cohort (n=91). Notably, because measured bulk gene expressions were unavailable in this cohort, Path2Omics [41] was applied to obtain inferred bulk gene expressions from whole slide images. Normalized LIME weights were used to demonstrate the contribution of each spatial cluster to the prediction of immunotherapy. (**j, k**) Comparison of spatial biomarkers compared to FDA-approved clinical biomarkers. The AUC value (**j**) and odds ratio (OR) on a log scale (**k**) achieved by our spatial-based biomarker (green) are shown, in comparison with Combined Positive Score (CPS; **left**, teal), Tumor Proportion Score (TPS, **middle**, light teal), and Combined Clinical score (derived by averaging all available CPS and/or TPS scores per patient; **right,** dark teal). Error bars denote 95% confidence intervals estimated from 1,000 bootstrap iterations. Statistical significance is indicated as *p≤0.05, **p≤0.01, and ***p≤0.001. Dashed vertical line indicates OR=1 (no effect).

Interpretation of model coefficients revealed that clusters 1 and 11 has the strongest association with HPV-positive tumors, sharing proliferative and immune-active transcriptional program, with antiviral immune landscape, consistent with the classically immunogenic microenvironment of HPV-positive HNSC(**Fig. 6c**). In contrast, clusters 3 and 5 exhibited negative coefficients favoring HPV-negative status, characterized by inflammatory, metabolic TME features(**Fig. 6c**). Together, these findings indicate that spatially resolved TME composition captures the distinct microenvironmental architecture underlying HPV-positive and HPV-negative HNSC and robustly stratify HPV status across geographically distinct cohorts.

### The Spatial TME cluster composition predicts patient response to concurrent chemoradiotherapy(CCRT)/radiotherapy and ICB

The clinical utility of spatially defined TME clusters was further evaluated by developing predictive models of response to cancer therapies. Specifically, we analyzed two independent patient cohorts: one treated with CCRT/radiotherapy (n=224, TCGA-HNSC therapy cohort) and another treated with ICB (n=91, Taiwan multi-center immunotherapy cohort, recently generated).

Like HPV status prediction, we represented each sample in a low-dimension 11-bit vector describing the fractional abundances of each of the 11 shared cluster compositions in its TME (denoting the fraction of the overall slide assigned to a given cluster). For each treatment cohort, we systematically evaluated four machine learning classifiers, including Logistic Regression (linear), Support Vector Classifier (SVC; kernel-based), Random Forest, and Gradient Boosting (tree-based ensembles). Models were trained independently for CCRT/radiotherapy and the unpublished ICB therapy cohorts using the 11-bit TME composition vector. For the CCRT/radiotherapy cohort, clinical variables (age, tumor stage, HPV status) were incorporated. Model performance was optimized through 5-fold stratified cross-validation. Notably, to avoid data leakage, the hyperparameter tuning was conducted on only the training subsets, while the test sets remained completely unseen (See **Methods** for details). We found that the Logistic Regression and SVC achieved the highest performance for the TCGA-HNSC CCRT/radiotherapy and Taiwan multi-center immunotherapy cohort, respectively, outperforming alternative classifiers (**Supplementary Data Fig. 4a, b**). Indeed, this model selection strategy ensured robust evaluation across diverse algorithmic assumptions while maintaining computational feasibility given our sample sizes.

Specifically, in the TCGA-HNSC CCRT/radiotherapy cohort, models based on spatial clusters derived from *HEiST*-inferred ST data achieved an AUC of 0.74 and AUPRC of 0.94 (85% response rate), significantly outperforming models based on the measured bulk RNA-seq expression (AUC=0.60, AUPRC=0.90, p<0.001), and markedly outperforming traditional direct image-based models (AUC=0.53, AUPRC=0.85, p<0.001), that predict response without inferring molecular data (**Fig. 6d**).

To assess clinical utility, we calculated the odds ratios (OR), measuring the proportion of true responders among those predicted as responders and comparing it with the response rate among those predicted as non-responders (see **Methods** for details). Once again, the spatial cluster model had a significantly higher OR of treatment response (OR=4.91, p=0.0003) compared to those stratified using bulk expression–based predictions (OR=3.34, p=0.01) or the direct H&E image-based model (OR=1.64, p=0.21; **Fig. 6e**).

To investigate biological programs underlying treatment response, we ranked the spatial clusters by their model correlation coefficients (**Fig. 6f**) and examined the pathway enrichment profiles of the top-ranked clusters. We found that resistance-associated clusters 1 and 4 exhibited tumor-intrinsic proliferative programs with inflammatory signaling indicative of immune evasion and therapeutic resistance. In contrast, responder-enriched clusters 10 and 7 showed immune activation signatures suggesting functional anti-tumor immunity. These results demonstrate that spatial compositional features capture clinically relevant biological information inaccessible through conventional bulk profiling or direct image-based approaches.

In the Taiwan multicenter immunotherapy cohort, the spatial cluster-based model achieved an AUC of 0.70, AUPRC of 0.51 (33% response rate), and ORs of 4.10. Once again, this significantly outperformed both Path2Omics [41]-inferred bulk expression-based (measured bulk expression unavailable) models (AUC=0.56, AUPRC=0.39, ORs =0.78, p=0.04) and direct image-based models (AUC=0.53, AUPRC=0.36, ORs=0.85, p=0.01; **Fig. 6g, h**). These consistent results across distinct therapeutic modalities demonstrate the superior predictive power of spatial representation compared to conventional biomarkers.

To enhance model interpretability, given the non-linear nature of SVC, we applied Local interpretable model-agnostic explanations (LIME) [42] to identify spatial features driving predictive performance and derive feature importance weights from the trained model (**Fig. 6i)**. This analysis uncovered two main resistance-associated clusters (2, 4) enriched for proliferative signatures, and two key response-associated clusters (6, 10) with activated immune activation programs. Notably, clusters 4 (resistance) and 10 (response) played a strong predictive performance in both CCRT/radiotherapy and ICB therapies. The consistency of these patterns across independent cohorts and different clinical endpoints demonstrates that spatial transcriptional programs identified from inferred spatial expression delineate clinically meaningful determinants of therapeutic outcomes.

Finally, the predictive performance of the spatial TME composition was benchmarked against FDA-approved ICB biomarkers combined positive score (CPS: n=25), tumor proportion score (TPS: n=47) [43], and a combined clinical biomarker score derived by averaging all available CPS and/or TPS scores per patient (Combined clinical: n=48) (**Fig. 6j, k**) in the Taiwan multicenter immunotherapy cohort. Notably, the TME cluster composition demonstrated consistently superior discriminative ability in all clinical response measures, with AUCs of 0.71, 0.70, and 0.70 in comparison with CPS (AUC=0.62; **Fig. 6j, left**), TPS (AUC=0.50; **Fig. 6j, middle**), and the combined clinical score (AUC=0.50; **Fig. 6j, right**), respectively. ORs analyses further reinforced these findings as the spatial biomarker-defined responders exhibited significantly higher odds of treatment response (OR=5.22–6.64; p=0.01–0.03), compared to the stratification by CPS (OR=4.47; p=0.138; **Fig. 6k, left**), TPS (OR=2.22; p=0.232; **Fig. 6k, middle**), or combined clinical score (OR=2.08; p=0.26; **Fig. 6k right**). Collectively, these findings establish the spatial cluster-composition-based spatial biomarker as a statistically robust predictor of ICB therapy response, outperforming all currently approved clinical biomarkers, and underscoring its translational potential for patient stratification.

## Discussion

We hypothesized, that a comprehensive understanding of the spatial heterogeneity and organization of the TME can considerably improve risk stratification and therapeutic decision-making in HNSC. To address this, we developed a clinically scalable framework, *HEiST*, that integrates routine H&E WSI with pathology foundation models to infer ST profiles in HNSC and discover spatially resolved biomarkers predictive of survival, HPV-status, and treatment response. To the best of our knowledge, this study presents the first ST inference and large-scale spatial analysis to systematically deconvolve and characterize the HNSC TME through its principal spatial constituents.

We demonstrated that *HEiST* robustly predicted spatial gene expression of approximately 2,800 genes from WSIs, as validated across two external ST cohorts, which are strongly enriched with known HNSC-related cancer hallmark pathways. By applying *HEiST* to the TCGA-HNSC cohort, we identified 11 reproducible spatial clusters that represent distinct clinically relevant TME compositions of immune, stromal, and tumor-associated programs within individual tumors. Unsupervised clustering of these spatial TME organizations revealed two robust subtypes of HNSC tumors — *Immune*-*Exhausted* and *Immune*-*Activated SpatioTypes*. These *SpatioTypes* were consistently observed across large independent cohorts, including the HANCOCK and independently collected Taiwan CCRT datasets, and demonstrated significant prognostic discrimination. Critically, spatial cluster composition accurately predicted HPV status, revealing a strong association between HPV-positive tumors and distinct spatial clusters. This finding has potential implications for understanding HPV-related immune evasion and therapeutic susceptibility in HNSC. Subsequently, spatial-cluster-based predictors significantly outperformed conventional bulk gene expression (both measured and slide-inferred) and direct H&E image-based (without spatial inference) approaches in predicting response to both CCRT/radiotherapy and prospectively collected ICB therapy cohort. Subsequently, interpretable modeling identified two specific clusters that consistently predicted resistance and response across both therapeutic modalities. This suggests that these spatial-molecular patterns may reflect fundamental determinants of therapeutic outcome across treatment contexts.

While several studies have applied H&E images to infer ST in large TCGA cohorts including breast [30], [31], kidney, colon, and ovarian cancer [31], these approaches have not been extended to HNSC with large-scale validation across multiple independent cohorts and varied treatment regimens. Importantly, the discovery of reproducible *SpatioTypes* specific to HNSC TME heterogeneity represents a distinct advance in HNSC biology. Their prognostic value was validated across three independent cohorts (TCGA-HNSC, HANCOCK, and the freshly acquired unpublished Taiwan CCRT). Further, we demonstrated their ability to infer the HPV status and predict patients’ response in two independent cohorts (TCGA-HNSC CCRT/radiotherapy, exclusively collected Taiwan multicenter immunotherapy), spanning different treatment paradigms. These results demonstrate the robustness and generalizability of our framework beyond single-institution or single-treatment contexts.

Our analysis has identified both shared and therapy-specific clusters associated with response to CCRT versus immunotherapy. Across modalities, Cluster 10 is broadly favorable, consistent with a permissive niche in which immune and tumor intrinsic programs are balanced and can support either genotoxic control or immune-mediated tumor elimination. Beyond the shared clusters, the dominant pro-response niche differs by therapy: CCRT benefit is linked to Cluster 7, marked by KRAS signaling up with MYC-associated proliferative activity, whereas immunotherapy benefit is driven by Cluster 6, which retains KRAS signaling but shows relative suppression of IL-6/JAK/STAT3 and TNF-α/NF-κB programs. Together, these patterns support a spatial framework with a shared permissive response niche, while the key determinants of benefit diverge between CCRT and immunotherapy through distinct microdomain-level biology. The clinical relevance of this spatial framework was further underscored by the robust predictive performance of the spatial biomarker relative to FDA-approved clinical biomarkers CPS, TPS, and their combination, supporting its translational potential for patient stratification in ICB therapy.

Despite its significant prognostic and predictive performance, this study has several limitations. Our discovery cohort, while diverse, represents a finite sample of HNSC spatial biology; model performance may vary in underrepresented tumor subtypes or anatomical subsites. Additionally, genes with low expression levels are challenging to predict accurately from histological features. Notably, the predictive power of individual genes depends on the variance of their expression across different spatial locations at TME. Moreover, class imbalance between responders and non-responders in both the CCRT/radiotherapy and ICB therapy cohorts could bias the predictive model toward the predominant group, thereby reducing its sensitivity and increasing the false negative rate. Furthermore, technical constraints of the 10X Visium platform, including limited spatial resolution and sparse gene capture, may limit the comprehensive characterization of the microenvironment. For instance, our discovery cohort (UoS) contains ∼ 18,000 expressed genes, whereas the external cohort UcL contains roughly half that number (∼ 10,500 expressed genes), which limits the scope of a comprehensive external validation.

Future work will focus on expanding training datasets to include more diverse anatomy and patient populations to improve robustness and generalizability of the underlying *HEiST* model. The predictive power of the model will be further enhanced by incorporating additional high-resolution spatial technologies, such as Visium HD and Xenium spatial single-cell platforms, to infer high-resolution microenvironmental patterns at the single-cell level. To further establish the clinical utility of spatially grounded response prediction models, retrospective external validation in even larger, multi-institutional cohorts will be performed, followed by prospective clinical trials.

In summary, this study presents a scalable approach for accelerating and democratizing precision oncology by identifying spatially grounded, cost-effective rapid biomarkers for survival and treatment response prediction in HNSC. This approach has already demonstrated its translational potential in breast cancer and now in HNSC, and we endeavor to extend it to other cancer types in the future.

## Methods

### Patients and tumor specimens

#### Spatial transcriptomics cohorts

We utilized one freshly generated and two publicly available spatial transcriptomics (ST) data from three independent cohorts. The discovery cohort comprised 10 slides (10 patients) from the University of Southampton (UoS) (7 HPV-positive, 3 HPV-negative) [11]. For external validation, we used two cohorts, 5 slides (5 HPV-negative patients) from the University of Hong Kong (HKU) [13] and 3 slides (3 HPV-negative patients) from UCLouvain (UcL) [12]. All cohorts included H&E-stained whole slide images (WSIs), spatially resolved transcriptomics data, and corresponding spatial coordinates.

#### TCGA-HNSC cohort

We studied publicly available 475 H&E-stained WSIs from 445 HNSC patients (324 male and 121 female, median [IQR] age: 61.0 [53.0–69.0]) from TCGA. Additionally, we collected patient-level clinical data, including demographics, pathological tumor stage, bulk RNA-sequencing data, and survival from the Genomic Data Commons. Besides, response to treatment outcome (patients received CCRT/radiotherapy) data were downloaded from UCSC Xena [44] for a subset of 224 patients (173 male and 51 female, median [IQR]: 61.0 [54.0–68.0]).

#### HANCOCK cohort

We included a total of 708 H&E-stained primary tumor WSIs from the HANCOCK dataset [34], comprising of 700 HNSC patients (558 male and 142 female, median [IQR] age: 61.0 [54.0–69.0]). Additionally, we retrieved demographics, tumor stage, and survival outcomes for all patients.

#### Taiwan CCRT cohort

We prospectively collected 252 H&E-stained WSIs of 213 HNSC patients (198 male and 15 female, median [IQR] age: 59.0 [52.0–66.0]) treated with concurrent chemoradiotherapy (CCRT) obtained from the National Science and Technology Council, Taiwan. Demographics, tumor stage, and survival outcomes for all patients were incorporated in the analysis (IRB No. 2021-04-004A).

#### Taiwan multicenter immunotherapy cohort

We analyzed a newly assembled cohort of 91 HNSC patients (94 H&E-stained WSIs) collected from the National Science and Technology Council, Taiwan (33 patients, 33 slides) and the Big Data Center of Taipei Veterans General Hospital, Taiwan (58 patients, 61 slides). Among these patients, 30 were immunotherapy responders, and 61 were non-responders. Patients received various treatment regimens, including ICB monotherapy, ICB with chemotherapy, ICB with targeted therapy, and triple combination therapy (IRB No. 2025-03-001CC).

### Spatial transcriptomics data preprocessing

#### Spot filtering

We retained spots with at least 25 expressed genes and with mitochondrial gene content below 20%.

#### Expressed gene filtering

We included genes expressed in at least 5% of spots within each sample. This resulted in 17,417 genes in the UoS training cohort and, for external validation cohorts, 13,166 genes in HKU and 10,879 genes in the UcL cohort.

#### Gene expression normalization

To stabilize variance and normalize expression distributions, gene expression values were log10-transformed with a pseudo count of 1.

#### Whole Slide Image preprocessing

We applied the same image preprocessing pipeline used in our previous works [46, 47]. Briefly, WSIs were divided into non-overlapping tiles, with tile size determined by the maximum distance between two consecutive spatial transcriptomics spots and resized to 224×224 pixels without spatial redundancy. We excluded tiles containing more than 50% background content. Finally, to reduce staining variation and batch effects, we applied Macenko’s method [47] for color normalization to the selected tiles.

### Spatial gene expression prediction framework

The framework consists of two main components:

#### Feature Extraction

To obtain robust tile-level image representations, we applied the pathology foundation model Virchow2. Virchow2 is a self-supervised vision transformer pretrained on 3.1M H&E-stained WSIs, which enables the extraction of rich, morphology-informed image embeddings. Each 224×224-pixel H&E-stained tile is encoded into a 2,560-dimensional feature vector that captures tissue morphology and cellular architecture.

#### Multilayer Perceptron (MLP) Layers

The Virchow2-encoded feature vectors were fed into multilayer perceptron (MLP) layers for spot-level gene expression prediction. The MLP comprised three fully connected layers with dimensions [2560 → 512 → 17,417] with ReLU activation functions between layers. Each output node represented the predicted expression level of a specific gene.

#### Model training procedures

The ST model was trained using the Adam optimizer (learning rate=0.0001), mean squared error as loss function, and a batch size of 32. To prevent overfitting and improve computational efficiency, we applied dropout regularization (rate=0.2) and early stopping with a patience of 50 epochs, respectively.

To ensure robust and unbiased model evaluation, we follow the leave-one-patient-out nested cross-validation approach as described in [45]. In each outer fold, one patient served as the test set; while the remaining 9 patients underwent 9-fold cross-validation (8 for training, 1 for validation) for model development. This patient-level splitting prevents information leakage across training, validation, and test sets.

For the UoS discovery cohort, predictions were obtained by averaging outputs from all inner fold models within each outer fold. Similarly, for external validation datasets (HKU and UcL) and clinical H&E cohorts (TCGA-HNSC, HANCOCK, and Taiwan) final predictions were estimated by averaging outputs from all models trained across the 10 outer folds.

#### Spatial Smoothing on Gene Expression

Following our previous approach [30], we applied spatial smoothing to reduce technical noise in ST data. Specifically, each spot’s gene expression was averaged with that of its nearest neighbors (approximately 8) within a 200-micrometer radius. This smoothing procedure was applied to both model-predicted and measured spatial gene expression during cross-validation within the UoS cohort, and during external validation in the HKU and UcL cohorts. For downstream analyses of patient survival and treatment response, smoothing was applied only to the model-predicted spatial gene expression, as measured spatial expression was not available.

Without smoothing, the model achieved a median correlation of 0.29 in cross-validation, with 5,248 robustly predicted genes (PCC>0.4) (**Supplementary Fig. 1a**), whereas external validation achieved a median of 0.132 with 445 high-correlation genes (**Supplementary Fig. 1d**). However, our primary goal was to apply the pre-trained model to large cohorts, lacking measured gene expression data for clinical applications. Our previous work in breast cancer [30] demonstrated that spatial smoothing enhances both molecular prediction accuracy and biological interpretability, making it well-suited for clinical outcome analysis.

#### Performance evaluation metrics

We evaluated model performance by Pearson Correlation Coefficient (PCC), a standard metric for evaluating ST prediction from histology images. PCC quantifies the linear relationship between predicted and measured values in terms of both strength and direction, defined as:

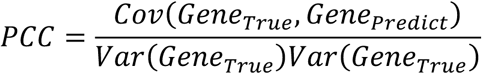

where *Cov* (*Gene_True_*, *Gene_Predict_*) is the covariance between the measured and predicted expression, and *Var* represents the respective standard deviations. For each patient, PCC was computed per gene across all spots, then aggregated across all genes to obtain a patient-level score. Overall model performance was reported as the median PCC across all patients.

#### Pathway enrichment analysis

We performed gene set enrichment analysis using the Enrichr algorithm implemented in the GSEAPY Python library (v1.1.9) with MSigDB Hallmarks gene sets and Benjamini-Hochberg false discovery rate (FDR) correction.

#### Spatial clusters

We implemented a two-stage clustering approach to identify uniform spatial clusters across samples. First, SpaGCN was applied to each TCGA-HNSC sample to identify distinct spatial domains based on inferred gene expression and spatial coordinates. To harmonize the cluster assignments across samples, gene expression within each SpaGCN-identified clusters were aggregated across all TCGA-HNSC slides to generate representative domain signatures. These signatures were log-normalized (*Seurat’s* log-normalization) [48], followed by dimensionality reduction and clustered using *Seurat’s* FindClusters function. The process yielded 11 stable consensus spatial clusters, and subsequently the original SpaGCN-identified cluster assigned to one of these 11 reference clusters that recapitulated coherent spatial tissue architecture across the TCGA-HNSC cohort.

These 11 reference clusters’ annotations were transferred to clinical cohorts (HANCOCK and Taiwan) using an anchor-based approach. SpaGCN-identified domain in each clinical sample was mapped to the reference TCGA-HNSC-derived clusters using *Seurat’s TransferData* function, which identifies mutual nearest neighbors (anchors) between the reference and query datasets, calculates similarity scores, and projects the reference embedding onto the query data. This anchor-based approach ensures consistent spatial cluster assignments across cohorts based on transcriptomic similarity.

#### Pathway activity quantification

To characterize the biology and pathway activity across the spatial clusters across multiple samples, we employed single-sample gene set enrichment analysis (ssGSEA) implemented in the GSEAPY Python library (v1.1.9). ssGSEA transforms inferred gene expression profiles into normalized enrichment scores (NES) based on the MSigDB Cancer Hallmark gene sets. For each sample, we calculated the mean NES across all spots within each spatial cluster, followed by z-score normalization. The z-scored values were then averaged across all samples to obtain final pathway enrichment profiles for each cluster.

To evaluate cross-cohort concordance of pathway enrichment scores between TCGA-HNSC and HANCOCK, we computed the Spearman rank correlation across all paired cluster–pathway entries using the SciPy Python library (v 1.15.3). Statistical significance was assessed using a two-sided permutation test.

### Discovering *SpatioTypes*

#### Spatial composition-based patient clustering

To recognize patient subgroups based on spatial composition, we performed unsupervised hierarchical clustering on the z-score–normalized patient spatial cluster composition matrix using seaborn’s (v0.13.2) clustermap function with Ward’s minimum variance method and Euclidean distance.

#### Optimal cluster number selection

To identify the optimal number of patient clusters and their stability, we computed silhouette scores for clusters k=2 to 10 using bootstrapping. For each k, the bootstrapped silhouette score was calculated by comparing the average intra-cluster distance to the nearest inter-cluster distance. The highest silhouette score (0.138) was achieved for k=2, as displayed in **Supplementary Fig. 2b** first panel, which indicates a well-separated and cohesive clustering structure. This solution was further validated using Davies-Bouldin (DB) and Calinski-Harabasz (CH) indices (**Supplementary Fig. 2b,** last two panels), which also identified k=2 as optimal, with lower DB and higher CH values confirming optimal cluster definition. All analyses were performed using scikit-learn (v1.7.1).

### Survival analysis

The prognostic implications of *SpatioTypes* were evaluated across different survival endpoints (OS, PFS, DFS, RFS) using Cox proportional hazards regression and Kaplan-Meier analysis implemented in the lifelines Python package (v0.30.0). Log-rank tests were used to assess the statistical significance of survival differences between *SpatioTypes*.

### Models for HPV status prediction

#### Spatial cluster model

To stratify HPV status in HNSC, we trained a Logistic Regression model on TCGA-HNSC cohorts using 11-dimensional spatial cluster compositions feature. The feature standardization strategy and hyperparameters of the model were optimized by standard procedures using scikit-learn’s (v1.7.1) within a nested 5×5 cross-validation framework. Notably, to avoid data leakage, the hyperparameter tuning was performed on training data only, while the test sets remained completely unseen. Model training performance was assessed on aggregated out-of-fold predictions and quantified by AUC. The generalizability of the model was further evaluated through external validation on the independent HANCOCK cohort by aggregating the predictions in all folds.

#### Direct model

As a baseline, we employed a direct model that stratifies HPV status response from H&E histology images without the intermediate gene expression inference. The model used the same Virchow2 feature extraction and MLP architecture with a classification layer. Each WSI was divided into non-overlapping 500×500-pixel tiles. Using a weakly-supervised learning approach [50, 51], the model was trained to predict HPV status from tile features using a 5×5 nested cross-validation strategy with patient-level splitting. Patient-level predictions were obtained by aggregating tile-level scores across each slide. The generalizability of the model was further evaluated through external validation on the independent HANCOCK cohort by aggregating the predictions in all folds. External validation was performed by applying the final models on the HANCOCK cohort, and performance metrics was computed over aggregated predictions from all folds.

#### Bulk transcriptomics model

For comparison, a Logistic Regression model was trained on measured bulk RNA-seq data from the TCGA-HNSC cohort. The top 100 genes with the highest variance associated with HPV status were selected using SelectKBest with f_classif criterion (ANOVA F-statistic). Gene expression features were standardized using z-score normalization (StandardScaler) before training the models. Feature selection and standardization were performed within each cross-validation fold to prevent data leakage.

### Models for treatment response prediction

#### Spatial cluster model

We trained four machine learning classifiers (Logistic Regression, Random Forest, Support Vector Classifier, Gradient Boosting) separately for the concurrent chemoradiotherapy/radiotherapy and ICB therapy cohorts to predict treatment response from spatial cluster proportions. For the TCGA-HNSC treatment cohort, models were trained on 11-dimensional spatial cluster features combined with clinical variables. For the immunotherapy cohort, models were trained on spatial cluster features alone. Feature standardization and hyperparameter optimization were performed following the same procedure as described for the HPV status stratification model, including nested 5×5 cross-validation with fold-wise scaling parameter estimation. Logistic Regression and SVC achieved optimal performance for concurrent chemoradiotherapy/ radiotherapy cohort and immunotherapy cohorts, respectively. Performance was evaluated using aggregated predictions across test folds, with metrics including AUC and odds ratios at Youden-optimized thresholds (bootstrap n=1,000 for confidence intervals).

#### Direct model

Using the same model architecture, training procedure, and evaluation strategy as described for the HPV status stratification direct model, treatment response was predicted directly from H&E histology images, bypassing intermediate gene expression inference. Patient-level scores in cross-validation were derived by aggregating tile-level predictions across all slides per patient.

#### Bulk transcriptomics model

We applied the pre-trained Path2Omics model [41], originally trained on TCGA-HNSC, to predict bulk gene expression directly from H&E slides in the immunotherapy treatment cohort (as measured expression not available).

To assess the predictive performance of bulk gene expression, we applied the same analytical framework as spatial cluster-based models. For the TCGA-HNSC treatment cohort, we used measured bulk RNA-seq expression data from TCGA, whereas for the immunotherapy cohort, we used Path2Omics-inferred gene expression data from the Taiwan multicenter immunotherapy cohort. In both cohorts, feature selection (100 genes) and standardization were performed applying the same procedure as described for the bulk transcriptomics HPV status classification model. The same four classifiers were trained using 5-fold stratified cross-validation with the same hyperparameter search grids as spatial models. Besides, clinical variables were included for the concurrent chemoradiotherapy/ radiotherapy cohort to ensure fair comparison with spatial-based models.

### Evaluation metrics

The performance of the HPV status stratification and treatment response prediction model was evaluated by measuring the area under the receiver operating characteristic (ROC) curve (AUC) using functions computed with functions implemented in Scikit-learn (v1.7.1) [51].

To estimate the odds ratio (OR), we first determined the optimal decision threshold exclusively from the training data using the Youden index derived from the ROC. The Youden index *J* is defined as 𝑌*_j_* = 𝑇*_PR_* − 𝐹*_PR_*, where *Tpr* and *Fpr* are the true and false positive rates respectively. The optimal threshold was then applied to the held-out test folds to generate binary predictions. ORs were computed from the pooled predictions. Besides, to ensure numerical stability in the presence of zero counts, we applied the Haldane–Anscombe correction (adding 0.5). The corrected *OR* is calculated as *OR* = 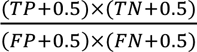, where *TP*, *TN*, *FP*, and *FN* represent true positives, true negatives, false positives, and false negatives, respectively. Finally, two-sided p-values and 95% confidence intervals were calculated under normal approximation.

### Statistics and reproducibility

We employed a rigorous train-validation-test split strategy with random assignment of samples from each data source to prevent data leakage and ensure unbiased performance evaluation. All evaluation metrics reported in this study were calculated exclusively on held-out test data that were never used during model development or hyperparameter tuning. No samples were excluded from any of the datasets. We quantified prediction uncertainty through bootstrap resampling of the training data, fitting multiple model instances to generate confidence intervals for all performance metrics.

Statistical comparisons between groups were performed using Spearman’s rank correlation for spatial cluster harmony. Survival analyses were conducted using Kaplan-Meier estimation with significance assessed by log-rank tests. Multivariable Cox proportional hazards models were used to adjust for clinical covariates (age and stage). All statistical tests were two-sided, with p<0.05 (∗p≤0.05, ∗∗p≤0.01, ∗∗∗p≤0.001) considered statistically significant unless otherwise specified. Multiple testing corrections were applied using Benjamini-Hochberg FDR where appropriate.

### Implementation details

The HEiST predictor was implemented in Python (v3.10.18) using PyTorch (v2.5.1) for deep learning and scikit-learn (v1.7.1) for machine learning analyses. Visualizations were generated using seaborn (v0.13.2) and matplotlib (v3.10.5) in Python, and ggplot2 (v3.5.1) in R (v4.3.2). Other analyses, including data processing and statistical analyses, were conducted using NumPy (v2.2.6), pandas (v2.3.2), OpenCV (v4.12.0), and OpenSlide (v1.3.1 for WSI processing), while pathway enrichment analysis was performed utilizing gseapy (v1.1.9). Survival analyses were performed using the lifelines package (v0.30.0).

## Authors’ contributions

S.B. developed the computational framework and performed all analyses. S.P. contributed to the development of machine learning models. S.P., S.R.D., A.S.,S.M, T.C, E.D.S., E.C., and C-P.D. interpreted the results and provided critical feedback on the study. G.J.T. and B.H.J provided the ST discovery dataset, G.J.T. and C.J.H provided critical feedback on the results. M.-H.Y., T.-H.C., S-K.T., and P-Y.C. contributed to the Taiwan CCRT and immunotherapy external validation cohort. T-H.C. managed clinical data collection, extraction, and curation, coordinated electronic file management, performed data cleaning and query resolution, and served as a liaison between clinical and research teams. S-K.T. and P-Y.C. facilitated the surgical procurement of tumor specimens and provided clinical annotation for all samples. Y.J.K and Y.C.Y provided pathological and clinical annotation for all samples. M-H.Y. served as collaborating PI, providing overall coordination, scientific input, and cross-team communication. D.-T.H. and E.R. supervised the study. S.B., D.-T.H., and E.R. wrote the manuscript with input and feedback from all co-authors.

## Supporting information

suplimentary_material_HEiST

## Acknowledgments

This research was supported by the Intramural Research Program of the NIH, NCI, Center for Cancer Research. The contributions of the NIH authors were made as part of their official duties, as NIH federal employees are in compliance with agency policy requirements, and are considered works of the US government. This work has utilized the computational resources of the NIH HPC Biowulf cluster (http://hpc.nih.gov) and Ceders-Sinai HPC cluster (https://intranet.cshs.org/sites/EISResearch/SitePages/High-performance-computing-(HPC).aspx). This research was also supported by Jim and Eleanor Randal Department of Surgery and Translational Research Institute of Ceders-Sinai.

## Declaration of generative AI and AI-assisted technologies in the writing process

During the preparation of this work, the author(s) used ChatGPT and Claude to improve the language and readability of the manuscript. After using these tools, the author(s) carefully reviewed and edited the content as needed and take full responsibility for the final content of the publication.

