## Supplementary material for "Histopathology-inferred spatial transcriptomics characterizes the tumor microenvironment in 1,500 head and neck tumors and predicts clinical outcomes": suplimentary_material_HEiST

### Supplementary Materials

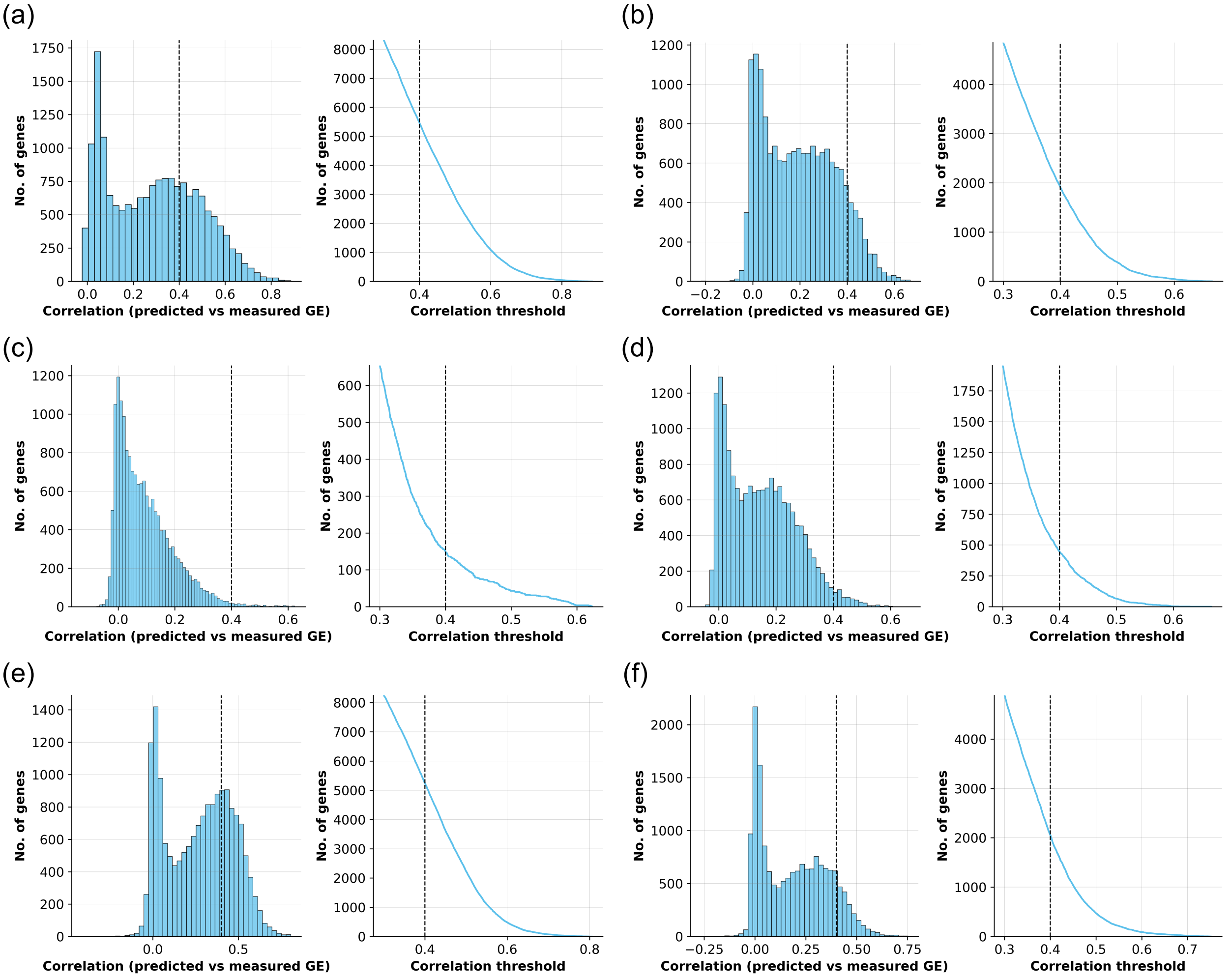


**Supplementary Fig. 1.** Evaluation of the spatial transcriptomics predictor. (a) Distributions of Pearson correlation coefficients (PCCs) and counts of robustly predicted genes across correlation thresholds without spatial smoothing, shown for cross-validation in the UniS cohort (dotted line, PCC = 0.4). (b–d) Prediction performance without smoothing in the UniHK (b), UCL (c), and combined independent validation cohorts (d). (e, f) Predictor performance with spatial smoothing applied in the UniHK (e) and UCL (f) cohorts.

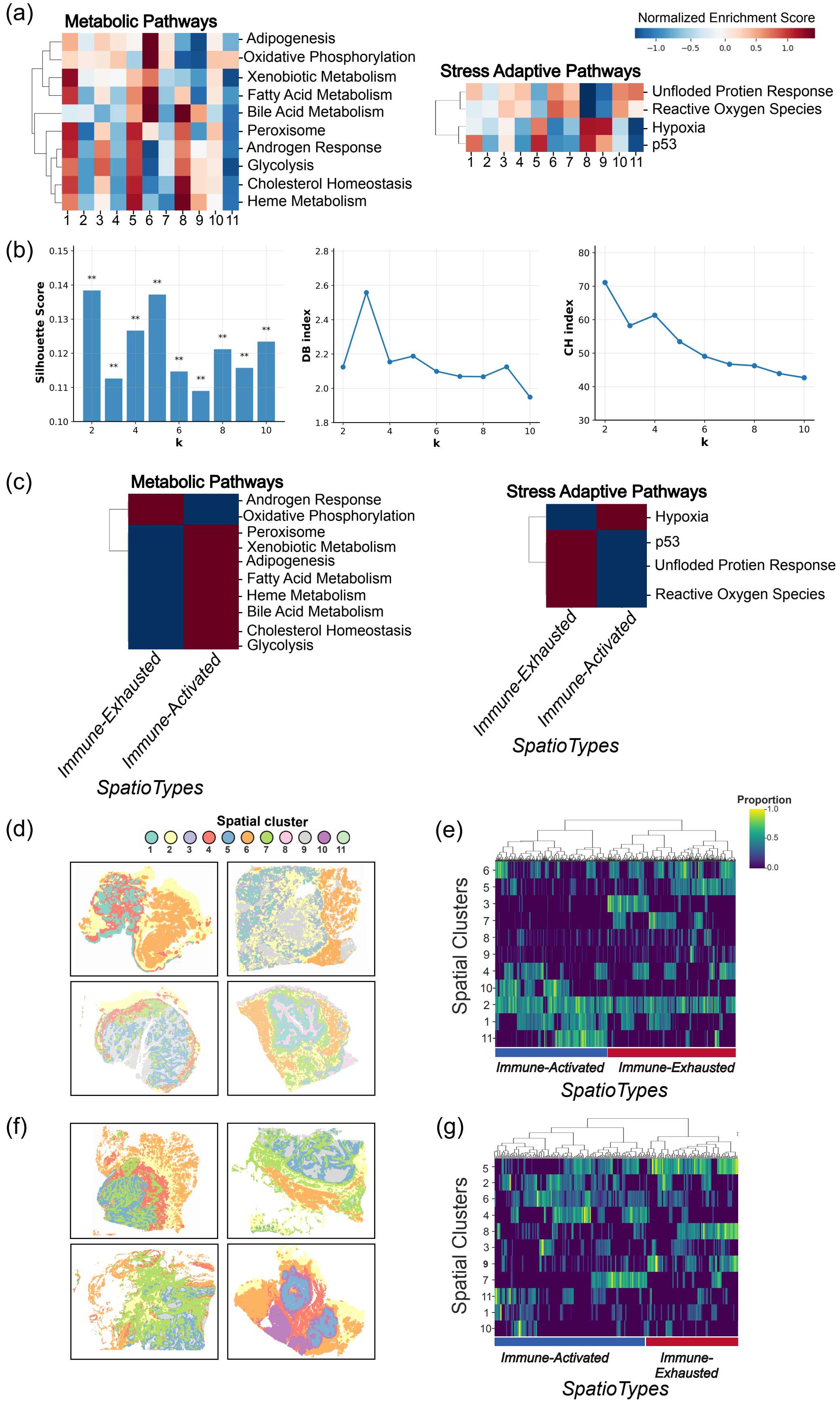
(a)
Metabolic Pathways
Adipogenesis
Oxidative Phosphorylation
Xenobiotic Metabolism
Fatty Acid Metabolism
Bile Acid Metabolism
Peroxisome
Androgen Response
Glycolysis
Cholesterol Homeostasis
Heme Metabolism
1 2 3 4 5 6 7 8 9 10 11
Normalized Enrichment Score
−1.0 −0.5 0.0 0.5 1.0
Stress Adaptive Pathways
Unfloded Protien Response
Reactive Oxygen Species
Hypoxia
p53
1 2 3 4 5 6 7 8 9 10 11
(b)
Silhouette Score
k
DB index
k
CH index
k
(c)
Metabolic Pathways
Androgen Response
Oxidative Phosphorylation
Peroxisome
Xenobiotic Metabolism
Adipogenesis
Fatty Acid Metabolism
Heme Metabolism
Bile Acid Metabolism
Cholesterol Homeostasis
Glycolysis
Immune-Exhausted
Immune-Activated
SpatioTypes
Stress Adaptive Pathways
Hypoxia
p53
Unfloded Protien Response
Reactive Oxygen Species
Immune-Exhausted
Immune-Activated
SpatioTypes
(d)
Spatial cluster
1 2 3 4 5 6 7 8 9 10 11
(e)
Proportion
Spatial Clusters
Immune-Activated
Immune-Exhausted
SpatioTypes
(f)
(g)
Spatial Clusters
Immune-Activated
Immune-Exhausted
SpatioTypes

**Supplementary Fig. 2.** Discovery of two prognostic *SpatioTypes*. (a) Pathway-level activity across the 11 spatial clusters shown as heatmaps of normalized enrichment scores for metabolic and stress-adaptive MSigDB Hallmark gene sets. (b) Clustering performance across k = 2–10 evaluated using the Silhouette score, Davies–Bouldin (DB) index, and Calinski–Harabasz (CH) index, with optimal values observed at k = 2. (c) Aggregated metabolic and stress-adaptive Hallmark pathway activity across the two *SpatioTypes*.(d, f) Representative tissue sections illustrating the spatial distribution of the 11 shared spatial clusters in the HANCOCK (d) and Taiwan CCRT (f) cohorts. (e, g) Unsupervised hierarchical clustering of patient-level spatial cluster proportions, with rows representing patients and columns representing spatial clusters, stratified by *SpatioTypes* in the HANCOCK (e) and Taiwan CCRT (g) cohorts.

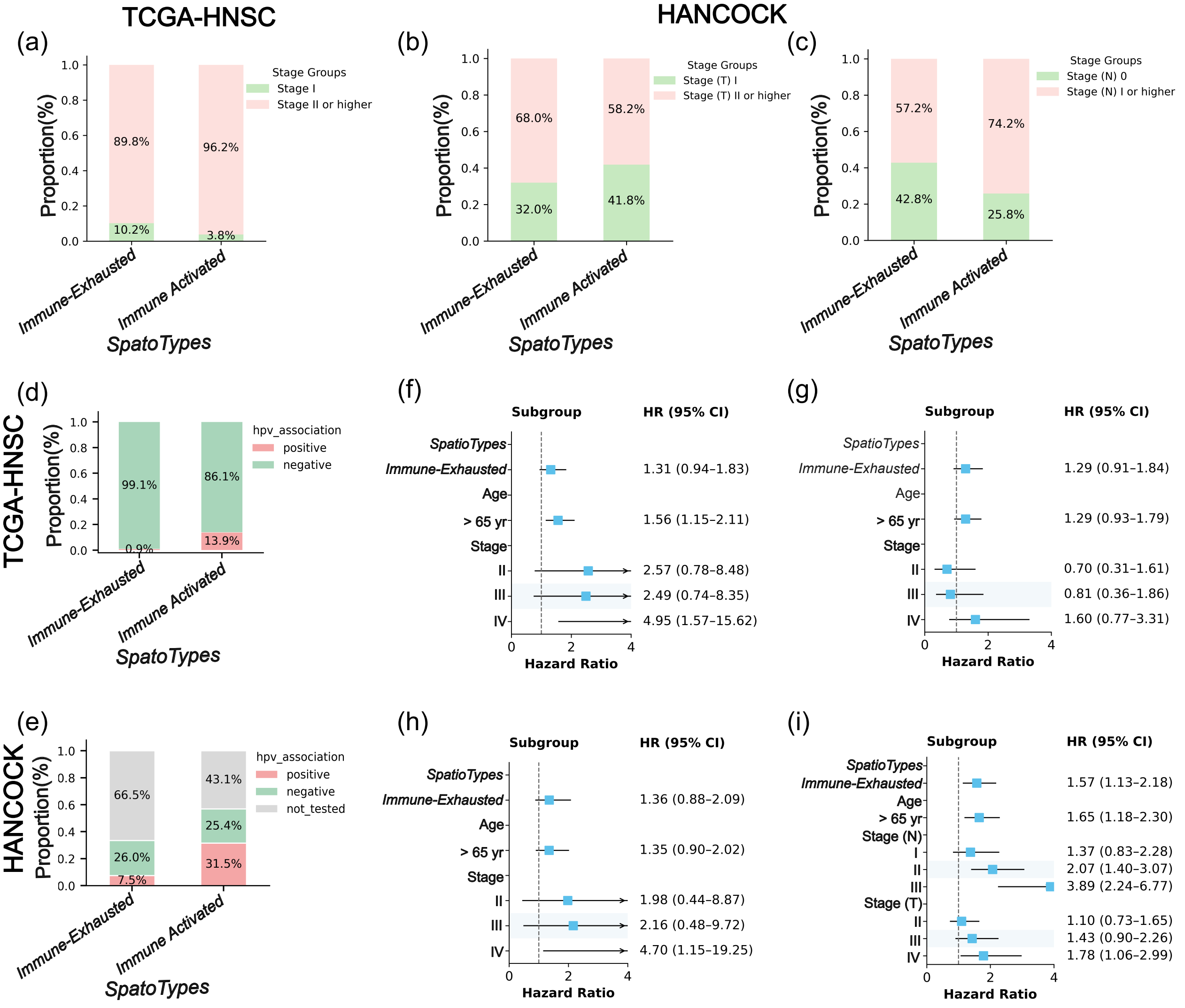


**Supplementary Fig. 3.** Clinical and molecular characteristics associated with *SpatioTypes*. (a–c) Distribution of clinical stage across tumor *SpatioTypes*, shown for overall stage for TCGA-HNSCC cohort (a), T (b), and N stage (c) for HANCOCK cohort. Bars represent the proportion of patients within each *SpatioType* stratified by stage group. (d, e) Distribution of HPV status across *SpatioTypes*, shown as proportions of HPV-positive and HPV-negative tumors in TCGA-HNSC (e) and HANCOCK cohort. (f-h) Forest plot of multivariate survival analysis for (f) overall survival, (g) progression free survival and (h) disease free survival in TCGA-HNSC. (i) Multivariable survival analysis for overall survival in the HANCOCK cohort. Forest plots show a consistent trend toward increased hazard associated with the immune-exhausted–like *SpatioType* across survival endpoints, independent of age and stage.

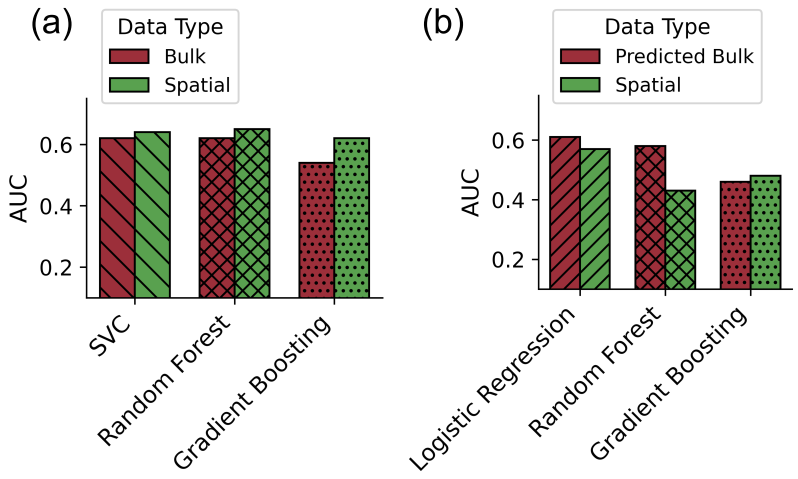


**Supplementary Fig 4.** Comparison of machine-learning models for predicting therapy response using bulk and spatial features. (a) Performance comparison of a Support Vector Classifier (SVC), Random Forest, and Gradient Boosting model for predicting adjuvant therapy response using measured bulk and spatial cluster–based features. (b) Performance comparison of Logistic Regression, Random Forest, and Gradient Boosting models for predicting immunotherapy response using predicted bulk and spatial cluster–based features. All models were hyperparameter-tuned using grid search with 5-fold cross-validation, and performance is reported as AUC.
